# Resolving clonal substructure from single cell genomic data using CopyKit

**DOI:** 10.1101/2022.03.09.483497

**Authors:** Darlan Conterno Minussi, Emi Sei, Junke Wang, Aislyn Schalck, Yun Yan, Alexander Davis, Hua-Jun Wu, Shanshan Bai, Cheng Peng, Min Hu, Anna Casasent, Alejandro Contreras, Hui Chen, David Hui, Senthil Damodaran, Mary E Edgerton, Scott Kopetz, Bora Lim, Nicholas Navin

## Abstract

High-throughput methods for single cell copy number sequencing have enabled the profiling of thousands of cells in parallel, yet there remains a significant bottleneck for data analysis. Here we present CopyKit, a comprehensive set of computational methods for the pre-processing and analysis of single cell copy number data to resolve clonal substructure and reconstruct genetic lineages in tumors. We performed single cell DNA sequencing of 2977 cells from multiple spatial regions in two liver metastasis and 7365 cells from three primary tumors with matched metastatic tissues. In the liver metastases, CopyKit resolved clonal substructure in different spatial regions, which revealed both clonal intermixing and spatial segregation in the tumor mass. In the matched metastatic colorectal and breast cancers, CopyKit resolved metastatic lineages and identified subclones and genomic events that were associated with metastases. These applications show that CopyKit is comprehensive tool for resolving copy number substructure in tumors.

## Introduction

Advances in single cell DNA sequencing (scDNA-seq) technologies for copy number profiling have led to vast decreases in costs and experimental worktime, while greatly improving the data quality and cell throughput. Techniques such as Direct Library Preparation (DLP)^1^, DLP+^2^, combinatorial indexing^3, 4^, Chromium Single Cell CNV^5^, and ACT^6^ have scaled-up copy number sequencing from dozens to thousands of cells, by using systems including nanowells, FACS and micro-droplet systems. These technologies have been applied to understand cancer evolution^7-9^, intratumor heterogeneity^2, 6^, DNA replication and cell cycle dynamics^2^, and normal tissue mosaicism^10, 11^. However, despite the rapid progress in technology development, there remains a significant bottleneck in the analysis of the resulting large-scale datasets that these platforms generate.

In contrast to single cell RNA-seq (scRNA-seq) technologies, where thousands of computational tools have been developed to pre-process and analyze data^12^, including Seurat^13^ and SingleCellExperiment^14^, there are limited user-friendly tools that provide complete workflows to analyze scDNA-seq datasets^15, 16^. In particular, the single cell genomics community still lacks a comprehensive package that can easily process and analyze scDNA-seq copy number data generated from sparse read count experiments. Currently, users face challenges in determining how to normalize read counts across the genome, identify the number of clusters that correspond to different biological subclones, and issue with comparing subclones to identify differences in their copy number profiles and their genetic relationships.

These challenges are important to address, since one of the most common application of scDNA-seq methods is to resolve intratumor heterogeneity (ITH) of copy number profiles within a patient’s tumor^1, 2, 9^. ITH is generated during the evolutionary expansion from a single somatic cell into billions of malignant cells, leading to different lineages and sets of chromosome aberrations within the same tumor mass^17, 18^. This ITH can be intermixed within the same spatial regions, or segregated across different areas of the tumor^19^. Additionally, chromosomal diversity can occur both at the primary tumor site, and in the matched metastasis, as specific subclones are often selected and expanded^20, 21^. Although methods including both deep-sequencing approaches and macro-spatial exome sequencing have been used to resolve subclonal differences within the tumor mass^22, 23^, single cell DNA sequencing (scDNA-seq) methods provide a more granular approach to resolve clonal substructure and understand the co-occurrence of genetics events in different subpopulations in the tumor mass^7, 8^. This information is important for understanding clonal dynamics during complex biological processes that occur during cancer progression, particularly in the context of therapeutic resistance, invasion, and metastasis^24-26^.

Here we present CopyKit, a comprehensive toolkit for the analysis of single cell copy number data that provides a suite of tools for performing binning of sequencing reads in genomic regions, copy number segmentation, quality control and downstream clustering and annotation analysis of scDNA-Seq datasets. We applied CopyKit to delineate the copy number substructure in different spatial regions from two liver metastasis and to study metastatic dissemination in matched samples from breast and colorectal cancer patients.

CopyKit R package can be installed from GitHub at **https://github.com/navinlabcode/copykit** and the complete documentation at **https://github.com/navinlabcode/CopyKit-UserGuide**.

### CopyKit Workflow

CopyKit extends the Bioconductor SingleCellExperiment^14^ class for the analysis of copy number datasets and is divided into four sequential modules: 1) Pre-processing, 2) Quality Control, 3) Data Analysis, and 4) Visualization. The pre-processing module starts from duplicate marked BAM files and generates count matrices by mapping the reads to variable genomic bins^9, 27^ (Fig. 1, step 1). Importantly, CopyKit’s requirement of duplicate marked BAM files is platform-agnostic. We provide assemblies for hg19 and hg38 at different variable genomic bin resolutions (∼55kb – 2.8Mb) that are user-selected. Additionally, user-defined scaffolds can be used for custom genome assemblies, genomic resolutions, or species. The binned counts are a mixture of Negative Binomial distributions with increased variance at higher counts. To address the overdispersion, CopyKit performs a variance stabilization transformation (VST) of the count matrix using the Freeman-Tukey transformation^28^ (Fig. 1, step 2, Extended Data Fig. 1). Following VST, the ratio matrices are computed by performing a sample-wise normalization of the variance stabilized bin counts by their mean. Therefore, a value of 1 corresponds to the average copy number of the sample, whereas higher values reflect amplified regions, and lower values represent genomic losses. Finally, a piecewise constant function is fit to the data using circular binary segmentation (CBS) ^29^ or multi-sample (cell) segmentation with multipcf^30^ (Fig. 1, step 3). The resulting segmented ratio means matrices are saved into the CopyKit object and the workflow continues with the quality control module.

**Figure 1:**
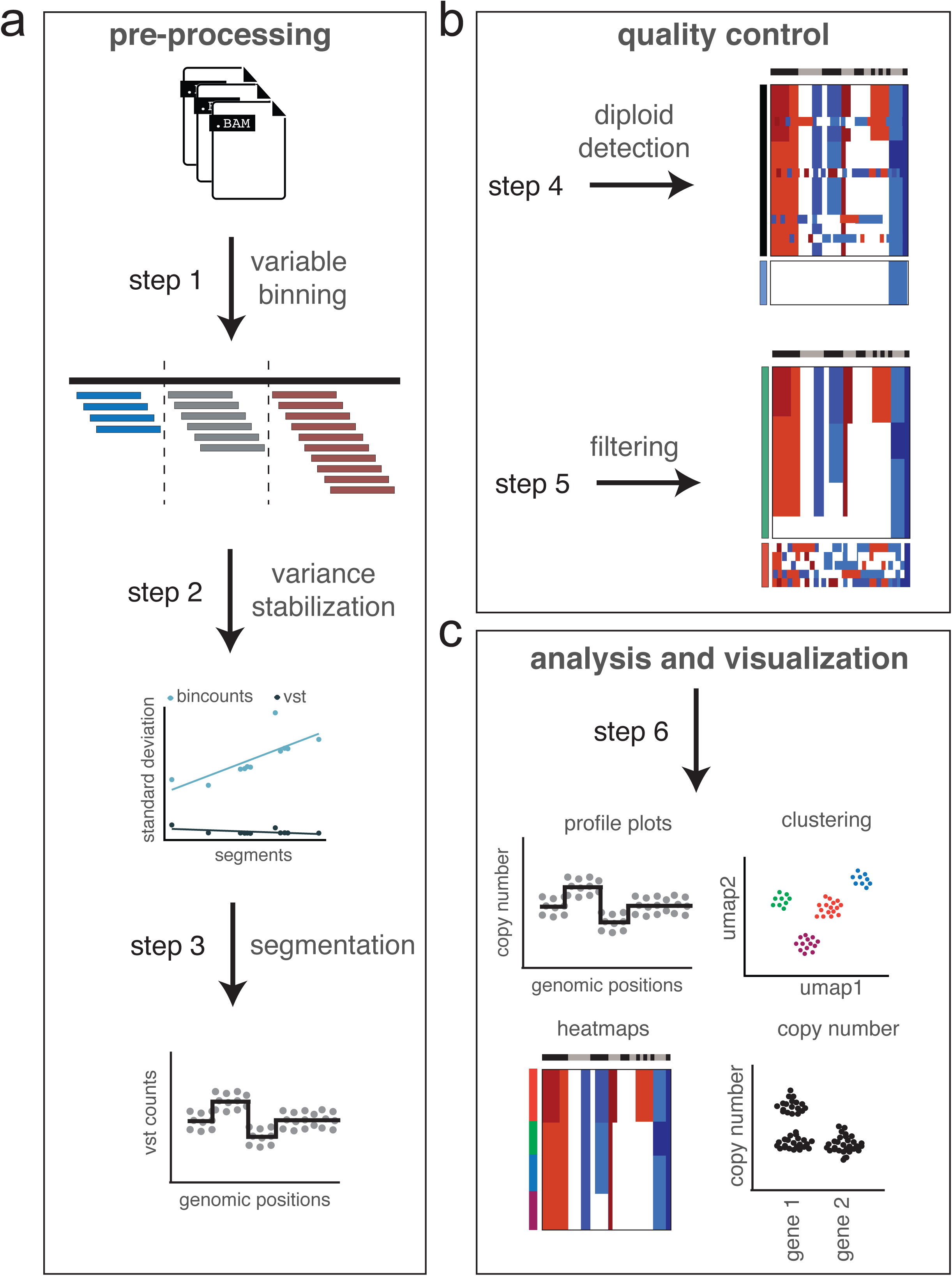
CopyKit Workflow and Data Analysis Modules. Schematic of the CopyKit workflow, which includes (a) data pre-processing, (b) quality control, and (c) analysis and visualization. The individual modules within each step include: variable bin read counting, variance stabilization, segmentation, detection of diploid cells, filtering of poor quality cells, data analysis and visualization. This workflow begins with BAM files of aligned sequence reads generated from a single cell DNA sequencing platform and ends with fully analyzed clustering and analysis of clonal genotypes from a tumor sample.

To retain high-quality aneuploid cells for downstream analysis we apply the CopyKit quality control (QC) module. Metrics from alignment and binning are stored in the CopyKit object metadata. To assess copy number data specific metrics, we calculate sample-wise mean overdispersion of the genomic bin counts, and count the number of chromosome breakpoints for each cell. We distinguish diploid from aneuploid cells by calculating the coefficient of variation (CV) of the segment ratio means for each cell. Next, we fit a gaussian mixture model to the observed CV distribution. The distribution with the smallest CV is assumed to be diploid. To increase the model sensitivity and prevent aneuploid cells from exclusion, we simulate the expected distribution of the CV for diploid cells (mean = 0, sd = 0.01). Using these criteria, we find that CopyKit can accurately detect diploid cells even with few cells present in the sample, as well as distinguish aneuploid cells with limited copy number events as small as one chromosome arm (Fig. 1, step 4, Extended Data Fig. 2, Methods).

CopyKit identifies low-quality cells by applying a nearest neighbor approach, using the correlation of the inferred copy number profiles. From the segment ratio means matrices, we calculate a Pearson correlation matrix across all samples. Next, we determine the sample-wise mean correlation to its k-nearest neighbors (default = 5) and compare it to a threshold (default = 0.9). Cells with a mean correlation smaller than the threshold are considered low-quality and, therefore, marked for removal from the dataset (Fig. 1, step 5, Methods). Altogether, the QC module ensures that high-quality cells are identified for downstream copy number analysis.

The CopyKit analysis and visualization modules work in synergy to study the scDNA-Seq datasets. To assess the copy number profile of individual cells, we provide an interactive app allowing the selection of cells. To identify subclonal populations, CopyKit leverages dimensionality reduction with Uniform Manifold Approximation and Projection (UMAP)^31^ as a pre-processing step. The reduced embedding is passed to density-based clustering methods (HDBSCAN)^2, 32^ or network-based (Louvain/Leiden)^33, 34^. A helper function performs a grid search and maximizes the Jaccard Similarity across parametrization values to evaluate clustering (Methods). Segment ratio means can be scaled with FACS derived ploidy values to obtain integer copy number states. To infer subclonal genotypes and lineages, we calculate consensus profiles across the single cells from each cluster (Methods), and compute distance-based phylogenetics with neighbor-joining^35^ or balanced minimal evolution^36^ at the single cell or consensus level. All analysis functions are accompanied by their visualization counterparts, with methods for plotting copy number ratio plots, UMAPs, heatmaps, consensus plots, gene-wise copy number states, and other data visualizations (Fig. 1, step 6, Documentation).

### Clonal substructure of two liver metastases

We flow-sorted nuclear suspensions by DNA ploidy distributions and performed scDNA-Seq using Acoustic Cell Tagmentation (ACT) of 2,977 cells from frozen liver metastasis of two different breast cancer patients (BL1, BL2). The first patient (BL1) was ER+/PR-/HER2-with an average ploidy of 4.30, while the second patient (BL2) was a triple-negative breast cancer (TNBC) patient that was ER-/PR-/HER2-with an average ploidy of 3.40 (Supplementary Table 1). We applied CopyKit to compute single cell copy number profiles with variable bins averaging ∼220kb resolution and analyzed the data to resolve clonal substructure. In sample BL1, CopyKit identified 607 diploid and 895 aneuploid cells (Fig. 2a). We excluded 102 cells marked as low-quality (resolution = 0.9) from the analysis (Fig. 2b). We assessed sample quality by calculating technical metrics (total read counts and PCR duplicates) and copy number intrinsic metrics (breakpoint counts and overdispersion). The BL1 cells averaged 1,035,783 total reads per cell (±246,883 s.d.) with a mean of 8.1% PCR duplicates (± 1% s.d.). We observe a significant difference of breakpoint counts (P-value = 1.87×10^−15^, Kruskal-Wallis test) and overdispersion (P-value = 0.047, Kruskal-Wallis test) between cells passing filtering QC compared to those marked for removal (Fig. 2c). Grid search of clustering parameters identified stability at k = 10, which was used for the clustering analysis (method = HDBSCAN). (Fig. 2d). The single cell data identified 4 subclonal populations (c1-c4) that shared a common evolutionary lineage (Fig. 2e, f).

**Figure 2:**
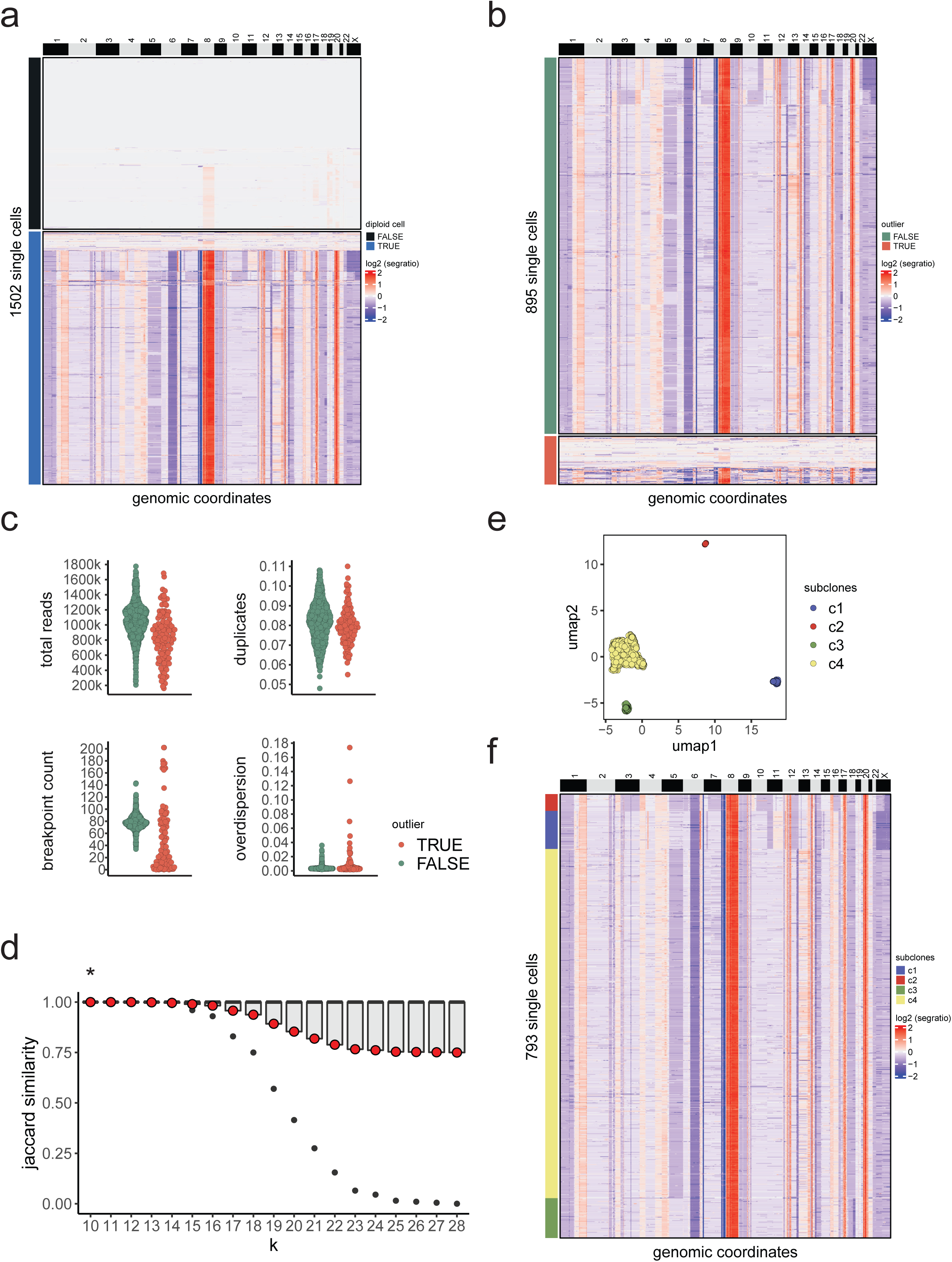
Example of the CopyKit workflow on a liver metastasis. **a**, Heatmap of single cell copy number profiles for the liver metastasis (BL1), in which diploid cells are identified and distinguished from the aneuploid cancer cells. **b**, Heatmap of single cell copy number profiles showing the identification and filtering of low QC cells from the dataset. **c**, Swarm plots of total number of reads and read duplicates (top) and overdispersion and counts of chromosome breakpoints (bottom), with colors representing the classification of cells by their data quality. **d**, Boxplot distribution of Jaccard Similarity from the clustering parameters grid-search for sample BL1 in which red dots represent the mean Jaccard Similarity and the star indicates the optimal K value selected for clustering. **e**, reduced dimension embedding UMAP and clustering of single cell copy number data in which colored points represents different clusters of subclones. **f**, Clustered heatmap of single cell copy number profiles for BL1, in which annotation bar represents classifications of single cells into subclones.

Similarly, in BL2, CopyKit identified 100 diploid and 1375 aneuploid cells with 108 cells marked as low-quality cells for removal (Extended Data Fig. 3a, b). The single cell data averaged 975,057 total reads per cell (± 303,911 s.d.) with a mean of 8.9% PCR duplicates (± 1% s.d.). We observe a significant difference between the total reads (P-value = 3.24×10^−13^, Kruskal-Wallis test) and overdispersion (P-value = 9.78×10^−9^, Kruskal-Wallis test) (Extended Data Fig. 3c). Jaccard similarity assessment identified k = 11 for clustering analysis (Extended Data Fig. 3d). Clustering identified 19 subclones with extensive copy number diversity within the tumor (Extended Data Fig. 3e, f) and marked 62 cells as outliers which were excluded from the analysis (Methods). These data show that CopyKit can efficiently remove low-quality cells and stromal/immune diploid cells from the downstream analysis and was able to resolve copy number substructure in two solid tumors.

### Spatial organization of subclones in two liver metastases

We further investigated the spatial organization of the subclones in BL1 and BL2 across different areas of the liver, since both samples were macrodissected prior to running scDNA-seq assays. In BL1 the 4 subclones across the 13 sections (S1-S13) showed both spatial segregation of subclones, and subclone intermixing in the same areas (Fig. 3a, b). Across all subclones (c1-c4) there were clonal CNA events including loss of 1p, 6q and 18, as well as gains of chromosome 8q (*MYC*) and 20 (*AURKA*). Sections S2 and S6-S7 had the highest Shannon diversity index (0.72, 95% CI [0.65, 0.81]) with mixtures of subclones c3/c4 (S2) and c1/c2/c3/c4 (S6-S7) in these regions (Fig. 3c). Section S3 was the only region that harbored a unique subclone (c4) compared to the other regions. Clone c4 was prevalent across the right side of the liver, whereas c2 prevailed across the left side (Fig. 3b). Subclone c2 showed large chromosomal losses on 3q, 7q, 10q (*PTEN*), and 16q (*WWOX*) and an amplification of chromosome 4p (*FGFR3*). Subclone c1 was restricted to sections S6-S7, showing a gain of chromosome 11q (*PGR*) and a focal amplification on 6q. Further, subclones c3 and c4 were spread across different liver sections and harbored losses of chromosome 5q (*APC*) in c4 and 18q (*BCL2*) in c3. Both subclones (c3 and c4) also had a gain of chromosome 13 (*FOXO1*) (Fig. 3d).

**Figure 3:**
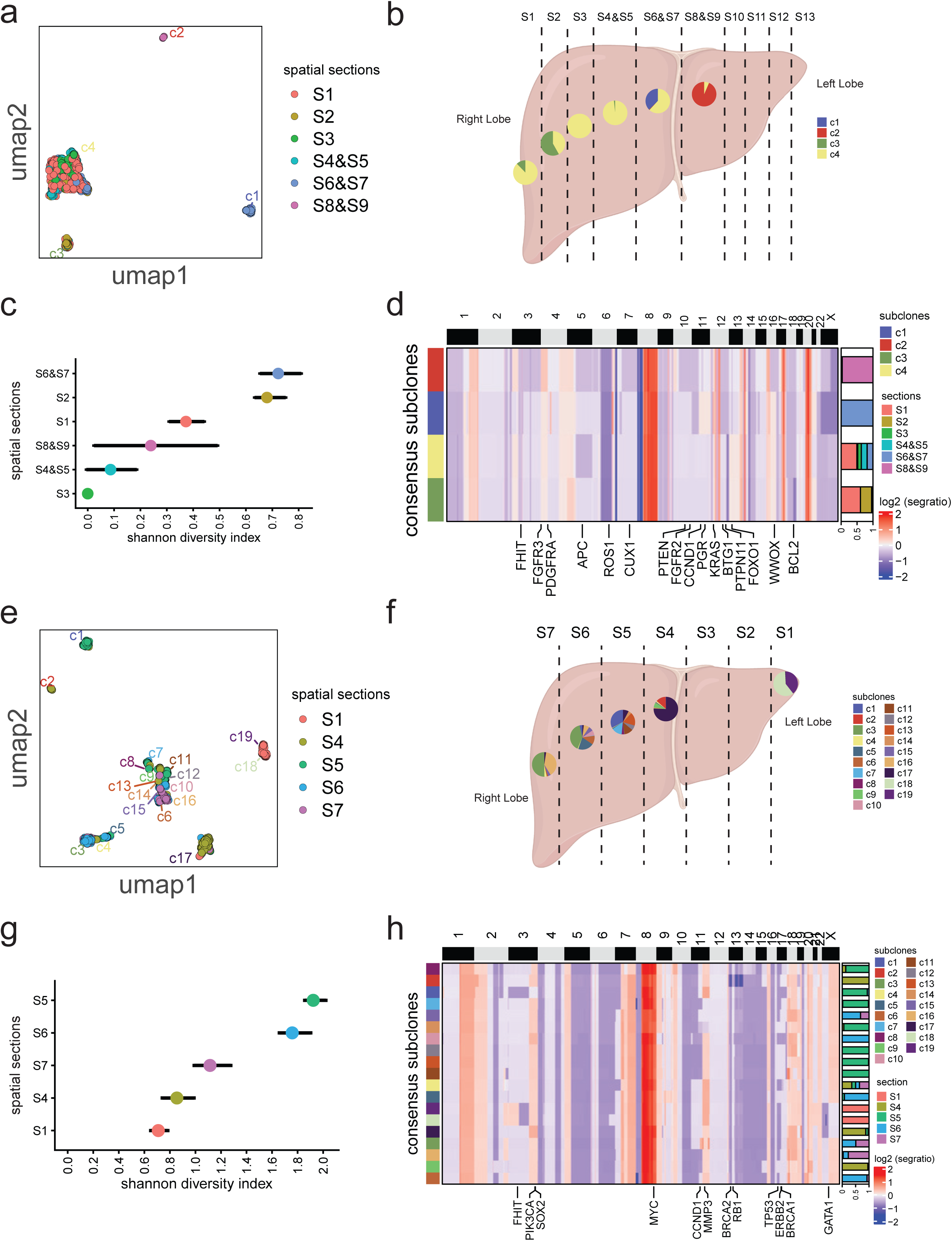
Spatial organization of subclones in two liver metastases. **a, e**, UMAP of single cell copy number data for the liver metastasis samples BL1 (a) and BL2 (e) in which colored points represents tumor sections. **b, f**, Schematic of liver macro-dissections of spatial sectors from sample BL1 (b) and BL2 (f), in which colored pie charts represent distribution of subclones across the sections. **c, g**, Shannon diversity indexes calculated from the single cell copy number data for BL1 (c) and BL2 (g) in which colored dots represent tumor sections and bars indicate 95% confidence intervals. **d, h**, Consensus heatmaps of subclones annotated with cancer genes, in which colored bar plots represent the frequency of a subclone according to the tumor’s spatial section (right side).

We further investigated the spatial organization of the 19 subclones in seven sectors (S1-S7) from the left to the right lobe of the liver from BL2. Similar to BL1, these data showed both spatial segregation of some subclones, and extensive subclone intermixing that varied across the tumor mass (Fig. 3e, f). Clonal CNA events across all the subclones included losses of chromosomes 5q, 6q, 13 and gains of 1q, 5p, 7q, 8q (*MYC*) 18 and 20 (*AURKA*). The spatial regions showed that S1, from the left side of the liver, had the lowest clonal diversity with a Shannon diversity index of 0.71 (95% CI [0.65, 0.78]), while the remaining sections (S4-S7) from the right side of the liver showed higher diversity indexes (range 0.85-1.75) with increased subpopulation numbers (range 7-11 subclones) (Fig. 3g). Subclone c2 was present only in section S4 with an additional chromosome 13 loss (*BRCA1* and *RB1*) relative to other subclones. In section S5, c1 was the predominant subclone with a loss of chromosome 3 and four subclones (c11, c12, c13, and c14) that shared a loss of chr11q (*CCND1*). Furthermore, subclones c8 and c9 were intermixed in sectors S4 and S5, while subclone c5 was shared between S5 and S6 and c7 was present across S4, S5 and S6. The subclone c17 displayed a gain of *SOX2* and a focal gain on chromosome 19 that was not detected in the other clones, and was the only subclone intermixed across all sectors, occupying the majority of sector S4. The c10 subclone was detected solely in section S6. The subclones c3, c6, c15, and c16, were intermixed in regions S6 and S7, together with subclone c4 which was also found in sectors S4 and S5. Finally, c18 and c19 were restricted to section S1, and showed exclusive gain of chromosome 3p and deletion of 3q and X compared to other clones (Fig. 3h). Overall, the CopyKit results showed examples of both spatial segregation of subclones, and extensive clonal intermixing, leading to different gains of oncogenes and losses of tumor suppressors across different areas of the tumor mass in both patients.

### Metastatic dissemination in a Breast Cancer Patient

To reconstruct clonal evolution across metastatic tissues, we performed single cell copy number profiling using ACT to profile 1521 cellsfrom a ER-positive, PR-positive, Her2-positive primary breast tumor (BM1). After quality-control and identification of diploid profiles we used 408 cells from the primary (n = 125) and its two matched metastatic sites: the liver (n = 238) and a pleural effusion sample (n = 45) (Supplementary Table 1). CopyKit identified four subclones, including two subclones in the liver metastasis that were distinguished by a gain on chromosome 5 (Fig. 4a-c). Phylogenetic analysis of the consensus subclones inferred a divergence of the primary tumor into two metastatic subpopulations to each organ site with subclone c1 from the liver metastasis presenting the shortest cophenetic distance to subclone c3 of the pleural metastasis (d=507.3, Manhattan distance) (Fig. 4d). The three tumors shared a common genomic lineage with clonal deletions on chromosomes 4p, 6q, 9p, 13, 22, and X, as well as gains of chromosome 12q and 16 (Fig. 4e). The lineage analysis further showed that the 17p (*ERBB2*) amplification, which was present in the primary tumor and was not detected at either of the metastatic sites, suggesting that this major subclone did not disseminate to the other organ sites. In contrast to the primary, the liver and pleural metastases had chromosomes losses of 18, 19, and 20p and a high-level focal amplification of a 6.8 Mb region on chromosome 8 that contained the cancer gene *FGFR1*. Furthermore, the liver metastasis showed two subclones (c1 and c2) distinguished by a gain on chromosome 5 (*FGF10* in c2), while the pleural effusion sample acquired additional focal events on chromosome 7 and 8, including gains of *BRAF* and *MYC*, as well as an amplification of chromosome 20 (*AURKA*) (Fig. 4d-f). Taken together, this analysis showed that after substantial copy number evolution at the primary site, a Her2-negative subclone from the primary breast tumor disseminated to the liver and acquired an *FGFR1* amplification and several chromosome losses, after which the cancer cells disseminated to the pleura (Fig. 4g).

**Figure 4:**
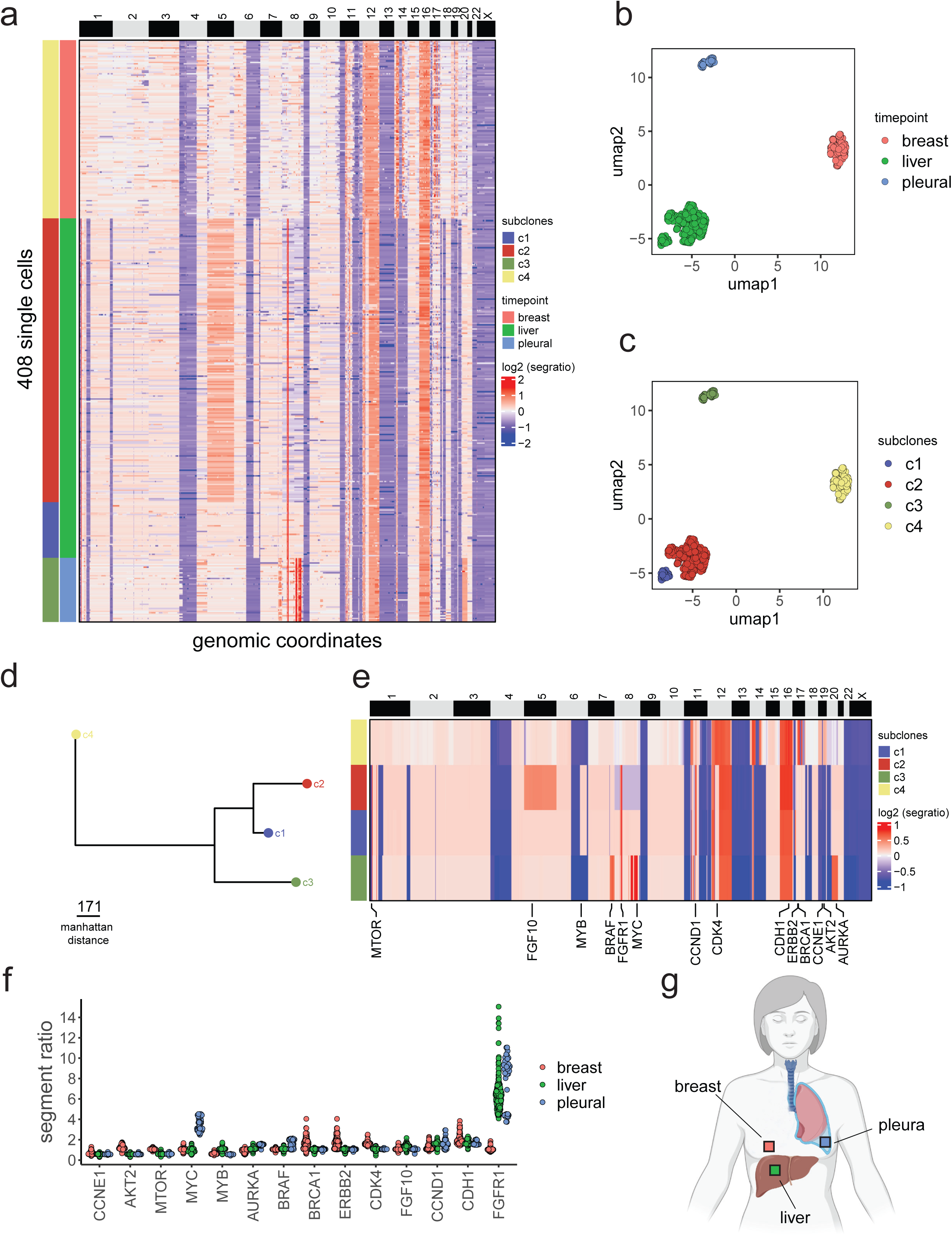
Dissemination of a primary breast tumor to two metastatic sites. **a**, Clustered heatmap of single cell copy number profiles from a primary breast tumor with matched metastatic liver and pleura sites, in which the left annotation bar represents the subclones and the right annotation bar represents organ site. **b, c**, Reduced dimension embedding UMAPs of single cell copy number data in which colored points represents organ sites (a) and subclones (b). **d**, Minimum evolution tree of consensus subclone genotypes for the different tumor sites. **e**, Consensus heatmap of subclones annotated with cancer genes. **f**, Swarm plot showing the distribution of inferred copy number events in which colored points represent the different tumor sites. **g**, Schematic of the organ sites where the tissues were collected.

### Metastasis of Colorectal Cancers to the Liver

We performed ACT of 5844 single cells and used CopyKit to analyze two primary colorectal adenocarcinomas (CM1 and CM2) and their matched liver metastatic samples. After quality control and identification of diploid cells we retained 4700 single cells for downstream analysis (Supplementary Table 1). In CM1, we identified 22 subclones across the primary colon (n = 1722) and liver metastasis (n = 1351), with no intermixing of subclonal genotypes across both organ sites (Fig. 5a-c). Clonal events across all subclones included losses of chromosomes 1p, while chromosome gains included 1q and 20, suggesting a common evolutionary lineage. Phylogenetic analysis of the consensus subclones revealed three main clades with a shared genomic lineage. Clade A1 reflected the primary colon subclones with lineage markers on chromosomes 1, 9, 13, and 15 that were shared by all subpopulations. The metastatic subclones from clade A2 diverged from the primary subclones with CNAs affecting several cancer genes including *SOX4, MYC, CDK8*, and *SMAD3*. Clade A2 in the liver metastasis was highly related to clade A1 in the primary tumor, with additional events chromosome 15, and showing a focal gain on chromosome 19, in addition to other subclonal events, suggesting a direct evolutionary lineage. However, the metastatic liver cells underwent additional copy number evolution forming clade A3 that diverged from the original metastatic population with gains of *PIK3CA* and deletions of *FHIT, CHEK1* and *IGFBP7* (Fig. 5d-f). These data show a clear progression in the metastatic lineage from clade A1 in the primary tumor to the clade A2 in the metastasis, which further expanded and diverged to form clade A3 in the liver mass.

**Figure 5:**
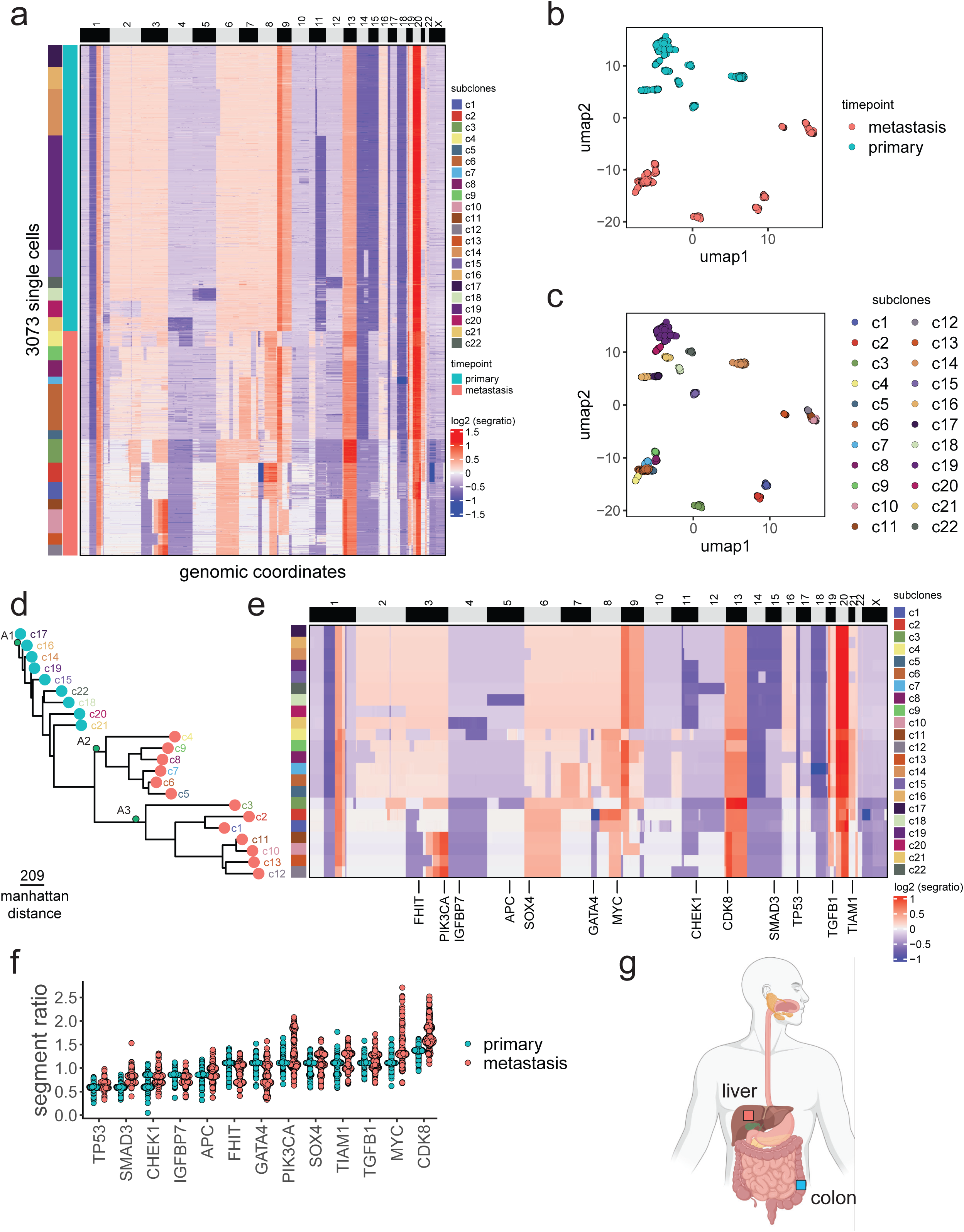
Dissemination from a primary colorectal tumor to a matched liver metastasis. **a**, Clustered heatmap of single cell copy number profiles from a patient CM1 with a primary colorectal tumor and matched metastatic liver tumor, in which the vertical left annotation bar represents subclones and the right annotation bar represents organ sites. **b, c**, Reduced dimension embedding UMAP of single cell copy number data, in which colored points represents tumor sites (a) and subclones (b). **d**, Minimum evolution tree of the consensus subclone copy number profiles, in which colored tip labels represent tumor sites and different clades of the tree (A1-A3) are labelled. **e**, Consensus heatmap of subclonal genotypes annotated with cancer genes. **f**, Swarm plot showing the distribution of inferred copy number events, in which colored points represent the different tumor sites. **g**, Anatomical schematic showing the organ sites from where the primary and metastatic tumor tissues were collected.

In the second patient (CM2), the CopyKit analysis of the primary colon (n = 897) and matched liver metastasis (n = 730) identified 8 subclones that shared a common evolutionary lineage as evidenced by multiple shared clonal CNAs, including gains of chromosomes 6, 7, 8q, 12p, 13, 16, X and losses of chromosome 1, 9, 11, 14. (Extended Data Fig. 4a-c). Phylogenetic analysis and clustered heatmap of the consensus subclones revealed three different clades during the divergence of the primary to the metastatic tumor. Clade A1 included an intermixed population (c6) with cells from both the primary and metastasis sharing the same genotype. The c6 subclone is very important, since it represents a pre-metastatic subpopulation in the primary tumor and constituted only a small percentage (6.6%) of the primary colon tumor mass. Clade A2 contained 5 subclones from the primary tumor (c8, c7, c9, c10). Clade A3 consisted metastatic liver subclones and included an amplification of chromosome 10 (*MAP3K8, FGFR2*) that was expanded at the metastatic liver tumor site (Extended Data Fig. 4d-f). Collectively, these data identified a pre-metastatic subclone in the primary tumor (c6) that disseminated and expanded at the metastatic liver site after acquiring an additional amplification of chromosome 10. Consistent with the CM1 patient, these data also show that most of the CNA events that were acquired during the expansion of the primary tumor were also represented at the metastatic liver tumor site.

### Application of CopyKit to Other scDNA-Seq platforms

We tested the application of CopyKit to data generated from two additional scDNA-seq platforms, DLP+ and Chromium Single Cell CNV (10X Genomics). The DLP+ dataset was generated from a normal lymphoblast cell line data that was previously published^2^. In total the CopyKit analysis of 6189 cells identified 552 outlier cells, and detected 5640 diploid cells with limited CNA events (Extended Data Fig. 5a). The QC metrics computed with CopyKit showed good performance in PCR duplicate rates and overdispersion metrics, with low chromosome breakpoints, as expected for normal diploid cells (Extended Data Fig. 5b). The Chromium 10X Genomics Single Cell CNV dataset was generated from a TNBC sample that was previously published (TN3)^6^. A total of 1185 single cells were analyzed with CopyKit, in which 18 cells were marked as diploid (Extended Data Fig. 6a) and 82 were marked as low-quality cells (Extended Data Fig. 6b). Analysis of the library metrics for total reads and PCR duplicates showed similar characteristics with no significant difference for cell quality (p-value = 0.351 and 0.098, Kruskal-Wallis test). The copy number specific metrics for counts of breakpoints and overdispersion detected significant differences between low and high-quality cells (p-value = 2.02×10^−7^ and 3.47×10^−8^) (Extended Data Fig. 6c). A Grid search of Jaccard Similarity selected k = 21 (Extended Data Fig. 6d) which was used for clustering analysis and revealed 5 outlier cells, that were excluded from analysis (Methods), and 7 subclones that shared a large number of clonal CNA events, but also harbored divergent CNA events (Extended Data Fig. 6e, f).

## Discussion

Here, we present a comprehensive toolkit called CopyKit for the pre-processing and analysis of scDNA-seq copy number data. CopyKit is agnostic to the technical platform, and is therefore broadly applicable to many scDNA-Seq technologies such as 10X CNV, DLP^1^, DLP+^2^, ACT^6^, and other future platforms that may be developed. We demonstrated applications of CopyKit in two important areas of cancer research: 1) resolving spatial subclonal structure in the tumor mass, and 2) reconstructing metastatic lineages. These data show that CopyKit is efficient at filtering and removing low-qualify and diploid cells, clustering subclones, computing consensus genotypes of subclones, annotating cancer genes and reconstructing clonal lineage trees.

In the two metastatic liver tumors, CopyKit revealed extensive subclonal structure indicating that copy number evolution continues at the metastatic tumors sites, well after seeding has occurred from the primary breast cancer. From the multi-regional sampling data, we identified both spatial segregation of subclones, and intermixing of genetically distinct subclones in the same macrodissected regions. The intermixing of subclones in the same spatial regions raise interesting questions about the possibility of subclonal interactions to support tumor progression, that have been described in other cancer types^9^ and murine systems^37^. Overall, these data are consistent with previous work on multi-region sequencing, showing distinct copy number events can occur across different sites during breast metastasis^38^.

We further used CopyKit to reconstruct metastatic lineages from the scDNA-seq data generated from matched primary and metastatic tumors of three cancer patients. In the breast cancer patient, we reconstructed a clonal lineage that showed a direct evolutionary trajectory from the primary breast tumors to the liver metastasis, which disseminated to the lung pleura and involved the acquisition of specific CNA events during this process, including a focal amplification of *FGFR1* that was present at both metastatic lesions. Most CNA events were already acquired in the primary breast tumor prior to the dissemination of the first subclones to the liver metastasis. Similarly, in the scDNA-seq data from both primary colorectal cancers and matched liver metastasis, CopyKit resolved the clonal substructure of the tumors and identified the subclones in the primary tumor that were most related to the metastatic genotypes. These data were consistent with a late-dissemination model of metastasis^21, 39^, by showing that most CNA events were acquired in the primary tumor prior to the dissemination and seeding of the metastases. Overall, these data show that CopyKit can resolve clonal substructure at both the primary and metastatic tumor sites, providing insights into the spatial organization of cancer cells and their clonal lineages during dissemination.

While there are many computational tools developed in R for copy number analysis of bulk DNA-seq data, they are generally limited in their scope and focus on individual processing tasks. For example, many methods have been developed for copy number segmentation from read count data^30, 40-43^. One tool that was developed for first-generation scDNA-seq data called Ginkgo^16^, a web interface that was designed for small scDNA-seq datasets and does not have advanced filtering and clustering tools that are necessary for processing large datasets from high-throughput technologies. Another tool is PhyliCS^15^, which introduced a python framework to perform filtering, clustering, and visualization of scDNA-Seq datasets, with a focus on quantifying spatial heterogeneity. However, pre-processing of the copy number dataset for segmentation, normalization and integer calls remains outside of the application programming interface (API). In comparison, CopyKit provides a comprehensive start-to-end framework to pre-process and analyze scDNA-seq copy number data, requiring minimal coding knowledge, with an API familiar to users of other single cell packages such as scRNA-Seq and analysis through Seurat^13^ or SingleCellExperiment^14^.

A notable limitation of CopyKit is the inability to resolve haplotype-specific copy number information and, therefore, cannot identify regions of copy number neutral loss of heterozygosity. This is in part due to the nature and sparseness of most scDNA-Seq data, and is not inherently a limitation to CopyKit. Some methods such as CHIESEL^44^ and schnapps^45^ have been developed and attempt to phase blocks of LOH across clusters of cells, but have limited genomic resolution. An important future direction of CopyKit is to develop multi-omics integration strategies (e.g. DNA&RNA, DNA&ATAC), to integrate data generated from different technologies, or analyze data from single cell co-assays that measure both modalities in the same cells, as newer technologies are developed.

In summary, CopyKit implements a comprehensive toolkit for the analysis of single cell copy number datasets, and addresses many aspects of an analysis workflow, from filtering to data visualization. We anticipate that CopyKit will reduce the analytical burden of scDNA-Seq copy number projects and will have broad applications for studying diverse areas of cancer biology, including intratumor heterogeneity, premalignant progression, metastasis, and drug resistance. CopyKit will also have utility for clinical applications in cancer diagnostics, treatment and drug target discovery. Beyond cancer, CopyKit may also have applications in pre-natal genetic diagnosis, neuroscience and understanding tissue mosaicism, or other areas where it is necessary to detect copy number events in single cells^46^. These studies will help elucidate the fundamental role of aneuploidy in tissues and the age-old question of whether aneuploidy is a cause or consequence of cancer progression.

## Data availability

Files used for this project are deposited to the sequence read archive (SRA) under BioProject PRJNA785341.

## Code availability

Code for the CopyKit R package and installation is available at: https://github.com/navinlabcode/copykit.

Complete documentation for CopyKit is available at: https://navinlabcode.github.io/CopyKit-UserGuide/.

Code to reproduce the figures from this manuscript is available at: https://github.com/navinlabcode/CopyKit_paper.

## Acknowledgments

We wish to express our gratitude to all the cancer patients and family members who kindly agreed to participate in this study and made this research possible. We are extremely grateful to Sandra Bishnoi and her family for supporting these research studies as well as the Advanced Breast Cancer (ABC) patient advocate group at MD Anderson. We would like to thank all of the cancer patients that have provided tumor specimens to support this work. We also thank the members of the MD Anderson Metastatic Breast Cancer Post-mortem Tissue Collection (PMTC) team, including Pheroze Tamboli, Michael Adams, Lashonda Johnson, Allison de la Rosa and Annie Wilson. This work was supported by grants to N.E.N. from the NIH National Cancer Institute (RO1CA240526, RO1CA236864), the Vivian Smith Foundation, the Emerson Collective and the CPRIT Single Cell Genomics Center (RP180684). N.E.N. is an AAAS Fellow, AAAS Wachtel Scholar, Damon-Runyon Rachleff Innovator, Andrew Sabin Fellow, and Jack & Beverly Randall Innovator. This study was supported by the MD Anderson Sequencing Core Facility Grant (CA016672). D.C.M is supported by a fellowship from the Schissler Foundation. We thank Andrew McPherson, Samuel Aparicio and Sohrab Shah for sharing the DLP+ dataset. We thank Michael Wigler, Jude Kendall and Michael Schatz for their work and contributions to early implementations of different software modules, including variable binning.

## Methods

### Human Tumor Samples

The human tumor samples collected from breast and colon cancer patients at MD Anderson were obtained under informed consent using two Institutional Review Board (IRB) approved protocols. All individuals consented to have their tissue used for research. The tissue samples were snap-frozen tissues that were dissociated into nuclear suspensions. A summary of the clinical tissue samples and metadata is provided in Supplementary Table 1.

### Isolation of single nuclei by FACS

Nuclear suspensions from the frozen tumor samples were prepared using NST-DAPI lysis buffer protocol as previously described^8^. Suspensions were filtered through a 40μm mesh prior to sorting. The DAPI intensity profile was used to gate the diploid and aneuploid population for all tumors. Single-nuclei were flow-sorted with BD FACSMelody or Beckman MoFlo Astrios into 384-well plates (Eppendorf 951020702). Depositing of single-nuclei was inspected under a microscope to ensure that single-nuclei were deposited to each well. Plates containing single-nuclei were spun at 1,500g for 5 min and sealed for subsequent processing.

### Acoustic Cell Tagmentation of single-nuclei

Acoustic Cell Tagmentation (ACT) was performed as previously described^6^. Briefly, FACS-sorted 384-well plates were spun at 1,500g for 5 min. Tagmentation reagents (Illumina FC-131-1096) were dispensed with the Echo525 system (Labcyte) and thoroughly mixed at every dispensing step. Nuclei were lysed in 200 nl (384PP_SPHigh) of freshly prepared Lysis buffer (Protease (1.36 AU ml^−1^) diluted 1:9 in 5% Tween 20, 0.5% Triton X-100, and 30 mM Tris pH 8.0) with the following thermocycler settings: 55 °C for 10 min, 75 °C for 15 min, and hold at 4 °C, lid temperature 80 °C and volume 1 μl. After lysis, 600 nl of tagmentation reaction mixture (TD:ATM 2:1, 384PP-Plus_GPSA) was dispensed with thermocycler settings: 55 °C for 5 min, hold at 4 °C, lid temperature 60 °C and volume 1 μl. The reaction was neutralized with 200 nl (384PP_SPHigh) of NT buffer for 5 min at room temperature. The final PCR reaction included 1.11uM N7XX and primers (384PP_AQBP) in 2X HiFi HotStart Ready Mix (Roche# KK2602, 6RES_GPSA) as previously described^6^. PCR reaction conditions were: 72 °C for 3 min, 98 °C for 30 s, (98 °C for 10 s, 63 °C for 30 s, 72 °C for 30 s) for 15–18 cycles, 72 °C for 5 min, hold at 4 °C, lid temperature 105 °C and volume 6 μl. ACT performance was evaluated by Qubit fluorometer and TapeStation (Agilent) from selected cell libraries. Final libraries were pooled and purified with 1.8X AMPURE XP beads. Libraries were sequenced at 50 single-end read cycles with dual barcodes on the Illumina HiSeq4000 or Illumina NextSeq2000 system.

### Demultiplexing and alignment

Sequencing reads were demultiplexed into single cell FASTQ files allowing 1 mismatch of the 8b barcode. FASTQ files were aligned to hg38 (GRCh38.p12) using bowtie2^47^ (v.2.4.4),converted from SAM to BAM and sorted with Samtools^48^ (v. 1.13). Duplicates were marked using Sambamba^49^ (v.0.70).

### Variable binning scaffolds for hg38 and hg19

Variable binning scaffolds (VarBin) for hg19 are available as previously described^9, 27^. Scaffolds for hg38 were created using Ginkgo^16^ pipeline following the protocol available at: https://github.com/robertaboukhalil/ginkgo/tree/master/genomes/scripts. Bins that intersected with ENCODE hg38 excluded regions^50^, centromere, and telomere regions were excluded with R package excluderanges^51^ (v.0.99.6). Additionally, 855 manually curated high-quality diploid cells were used to detect outlier bins. Bins that were found to be 2.5 s.d. above the mean bin counts in more than 1 cell were excluded from the scaffold. Chromosomes X and Y were excluded from the analysis.

### Inference of copy number

Sequencing reads from BAM files with marked duplicates are counted into VarBin assembly scaffolds using Rsubread^52^ (v.2.8.1). Chromosome Y was excluded from this analysis in all samples. BAM files with less than 10 mean bin counts were excluded from the analysis. Bin counts are normalized for GC content with Locally Weighted Scatterplot Smoothing (LOWESS) regression. GC-corrected bin counts are normalized using the Freeman-Tukey variance stabilization transformation^28^, let X be the GC-corrected bin count and Y be the variance stabilized bin counts:

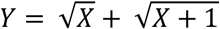

Sample-wise ratios are computed by taking the bin-wise ratio of the bin counts to the sample mean bin count. Segmentation was performed using CopyKit default parameter ‘CBS’ that uses circular binary segmentation (alpha = 1e^-10^, undo.splits = ‘prune’) from the Bioconductor DNAcopy package^41^ (v.1.68.0) followed by merge levels to join adjacent segments with non-significant differences (pv.thres = 1e^-10^) using package aCGH^53^ (v.1.72.0).

### Detection of diploid cells

To detect diploid cells CopyKit calculates the sample-wise coefficient of variation from the segment ratio means. The expected coefficient of variation for diploid cells *N*(0, 0.01) is simulated for *x* cells, where *x* contains an equal number of data points to the number of cells present in the dataset. An expectation-maximization algorithm is used to fit a mixture of gaussian distributions to the coefficient of variation from the samples together with the simulated dataset using R package mixtools^54^ (v.1.2.0). The distribution containing the simulated dataset is inferred to be the diploid distribution. Samples that group with the inferred diploid distribution and present coefficient of variation smaller than 5 standard deviations from the mean diploid distributions are classified as diploid samples. Single cells marked as diploid in tumor samples were excluded from downstream analysis. To assess precision and recall we simulated diploid single cells (n = 50) at 220kb resolution from a Poisson distribution (lambda = 50). To simulate aneuploid single cells, we added whole arm or whole chromosome gains (lambda = 100) and deletions (lambda = 25) from a Poisson distribution to the simulated cells. Diploid cells were added to the simulated aneuploid dataset incrementally from 1 to 50 single cells, replacing aneuploid cells.

### Filtering of low-quality copy number profiles

To detect low-quality cells, CopyKit calculates the Pearson correlation matrix of all samples from the segment ratio means. Next, we calculate a sample-wise mean of the correlation between a sample and its k-nearest-neighbors (default = 5). Samples in which the correlation value is lower than the defined threshold are classified as low-quality cells (default = 0.9). Samples were processed with default CopyKit settings with the exception of the BM1 primary breast sample, in which resolution was set to 0.8. Samples marked as low-quality were excluded from this work analysis.

### Calculation of Shannon diversity indexes

We calculated the Shannon diversity index by measuring the proportion of subclones for each section of the tumor as follows:

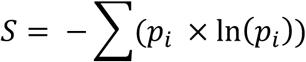

With 95% confidence intervals calculated by bootstrapping (R = 2000).

### Clustering parametrization

To determine cluster stability, CopyKit evaluates the Jaccard Similarity using R package ‘fpc’^55^ (v2.2-9) over a range from k=10 to the square root of the number of cells. The value that maximizes the Jaccard Similarity (default = ‘median’) is returned as the suggested value to be passed on for sample clustering algorithms.

### Clustering of samples

To identify subclonal populations the segment ratio means are embedded into a reduced dimensional embedding using Unifold Manifold Approximation and Projection (UMAP)^31^ with R package ‘uwot’ (v.0.1.10) (min_dist = 0, n_neighbors = 50, seed = 17). The resulting embedding is passed on to a clustering algorithm (default = ‘hdbscan’)^32^. Samples classified as outliers by hdbscan (cluster ‘c0’) were excluded from this work analysis. Samples were processed with default settings with the exception of sample BM1 (min_dist = 0.1, n_neighbors = 10). BM1 presented tumor-diploid doublet cells cluster that was excluded from the analysis (n = 24). BM1 cells were re-clustered after doublet exclusion.

### Calculating consensus profiles

To generate consensus profiles, we calculate the cluster-wise median of the *i*^th^ segment from samples assigned to the same cluster group.

### Phylogenetic reconstruction of consensus subclone trees

Pairwise distance matrices of consensus profiles were calculated using Manhattan distance. Phylogenetic inference were constructed using the balanced minimum evolution algorithm^36^ with R package ape^56^ (v.5.3). Trees were rooted with inferred MRCA profiles from primary samples (see ‘Inference of the most recent common ancestral’) and plotted using R package ggtree^57^ (v.3.2.1).

### Inference of the most recent common ancestor

To infer the most recent common ancestor (MRCA), among the consensus clusters from the primary sample, we selected the segment ratio mean value closest (L1 norm) to a the mean ratio value (default = 1).

### Processing of DLP+ datasets

Fastq files from sample SA928 A090533C^2^ were aligned and processed with CopyKit as previously described (see ‘Demultiplexing and alignment’ and ‘Inference of copy number’).

### Processing of Chromium 10X Genomics Single Cell CNV datasets

Fastq files from the Chromium Single Cell CNV (10x Genomics) experiments were processed as previously described^6^. Analysis was performed with CopyKit (see ‘‘Inference of copy number’, ‘Detection of diploid cells’, ‘Filtering of low-quality copy number profiles’, and ‘Clustering of samples’) with default settings, with the exception of UMAP embedding in which min_dist was set to 0.1).

### Parallelization

Whenever possible, CopyKit adopts the BiocParallel (v.1.28.0)^58^ framework for parallel computations to improve processing speed and timeframe.

### Statistical Analysis

Statistical analysis for this manuscript were performed in R software^59^ (v.4.1.1) with RStudio IDE^60^ (v.1.4.1717) and R package Rstatix^61^ (v0.7.0).

### Plots

Plots in CopyKit are generated using ggplot2^62^ (v.3.3.5) and ComplexHeatmap^63^ (v.2.10.0).

### Additional packages

Additional R packages used within CopyKit are: viridis^64^ (v0.6.2). GenomicRanges^65^ (v.1.46.0), tidyverse^66^ (v.1.31.0), scran^67^ (v.1.22.0), SummarizedExperiment^68^ (v.1.24.0), S4Vectors^69^, ggbeeswarm^70^ (v0.6.0), ggalluvial^71^ (v.0.12.3).

### Illustrations of Patient Anatomy

Figures 4g, 5g, and Extended Data Fig. 2g were created with BioRender.com

