## supplementary figures for "Resolving clonal substructure from single cell genomic data using CopyKit"

Minussi *et al., 2022*

### Extended Data

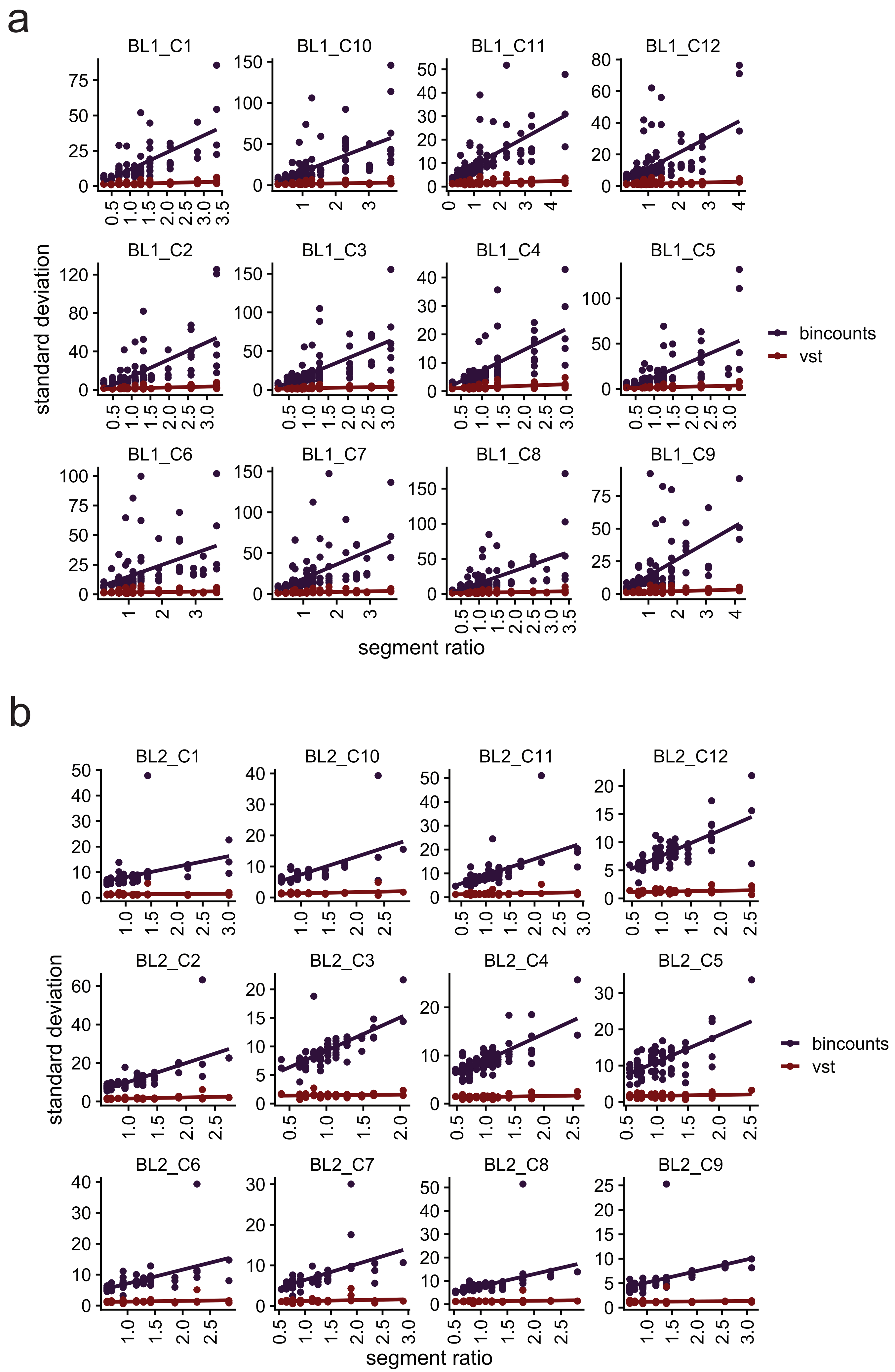
a
BL1_C1
BL1_C10
BL1_C11
BL1_C12
BL1_C2
BL1_C3
BL1_C4
BL1_C5
BL1_C6
BL1_C7
BL1_C8
BL1_C9
standard deviation
segment ratio
bincounts
vst
b
BL2_C1
BL2_C10
BL2_C11
BL2_C12
BL2_C2
BL2_C3
BL2_C4
BL2_C5
BL2_C6
BL2_C7
BL2_C8
BL2_C9
standard deviation
segment ratio
bincounts
vst

**Extended Data Fig. 1: Variance Stabilization Transformation.**
**a, b,** Scatter plot of standard deviation by segment mean ratios for 12 randomly sampled single cells from the BL1 and BL2 liver tumors with colors representing raw bin count values (blue) or Freeman-Tukey variance stabilized bin counts (red).

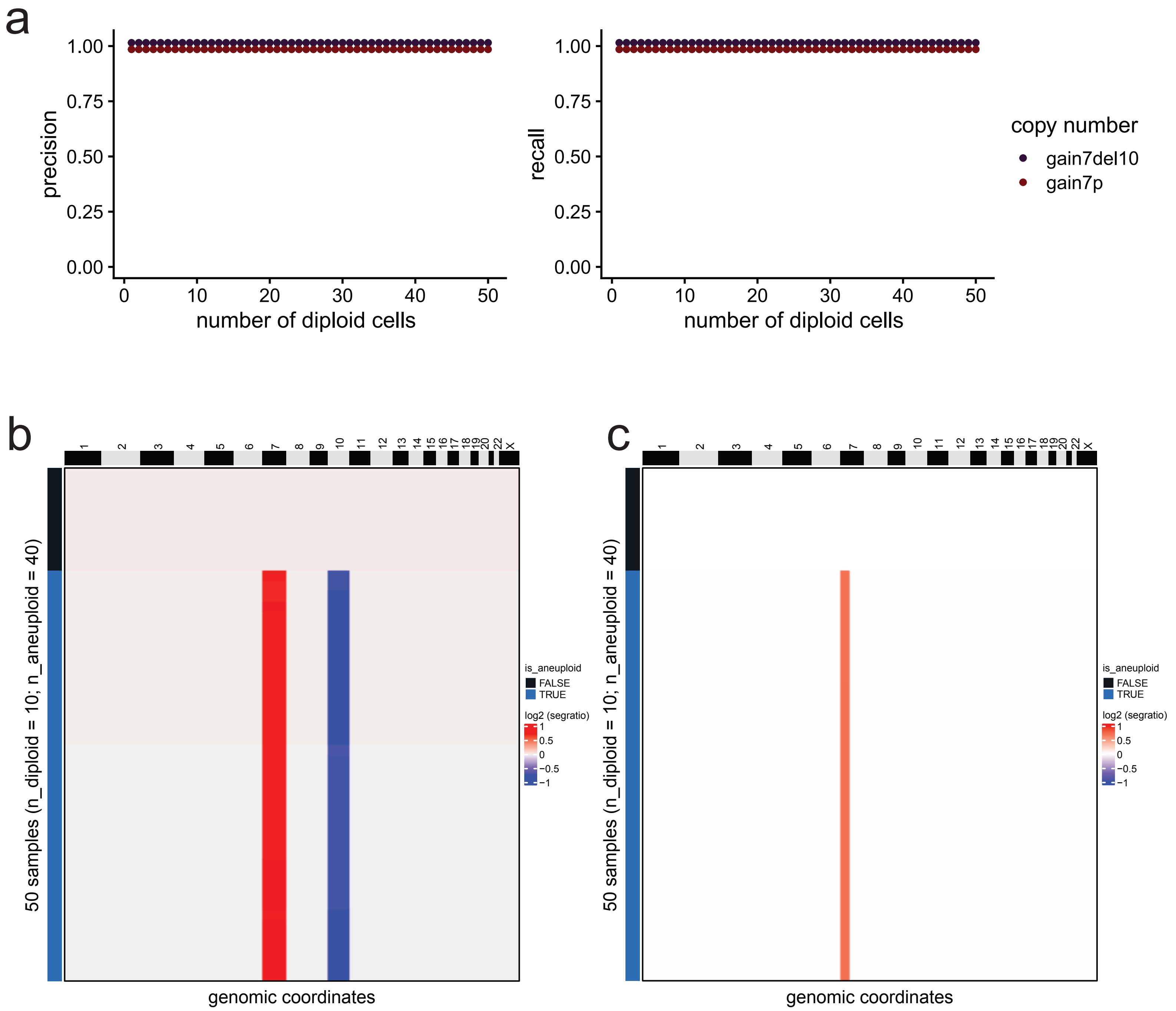

a
precision
recall
number of diploid cells
copy number
gain7del10
gain7p
b
c
50 samples (n_diploid = 10; n_aneuploid = 40)
is_aneuploid
FALSE
TRUE
log2 (segratio)
genomic coordinates

**Extended Data Fig. 2: Detection of diploid and aneuploid cells in simulated datasets.**
**a,** Precision and recall values for two 220kb genomic resolution simulated datasets (n = 50) with incrementally added diploid cells from 1 to 50 cells. **b, c,** Example heatmaps of 220kb simulated datasets with chromosome 7 gain and chromosome 10 deletion (a) and chromosome 7p gain (b), in which annotation bars show the classification of cells into diploid (n = 10) or aneuploid (n = 40) groups.

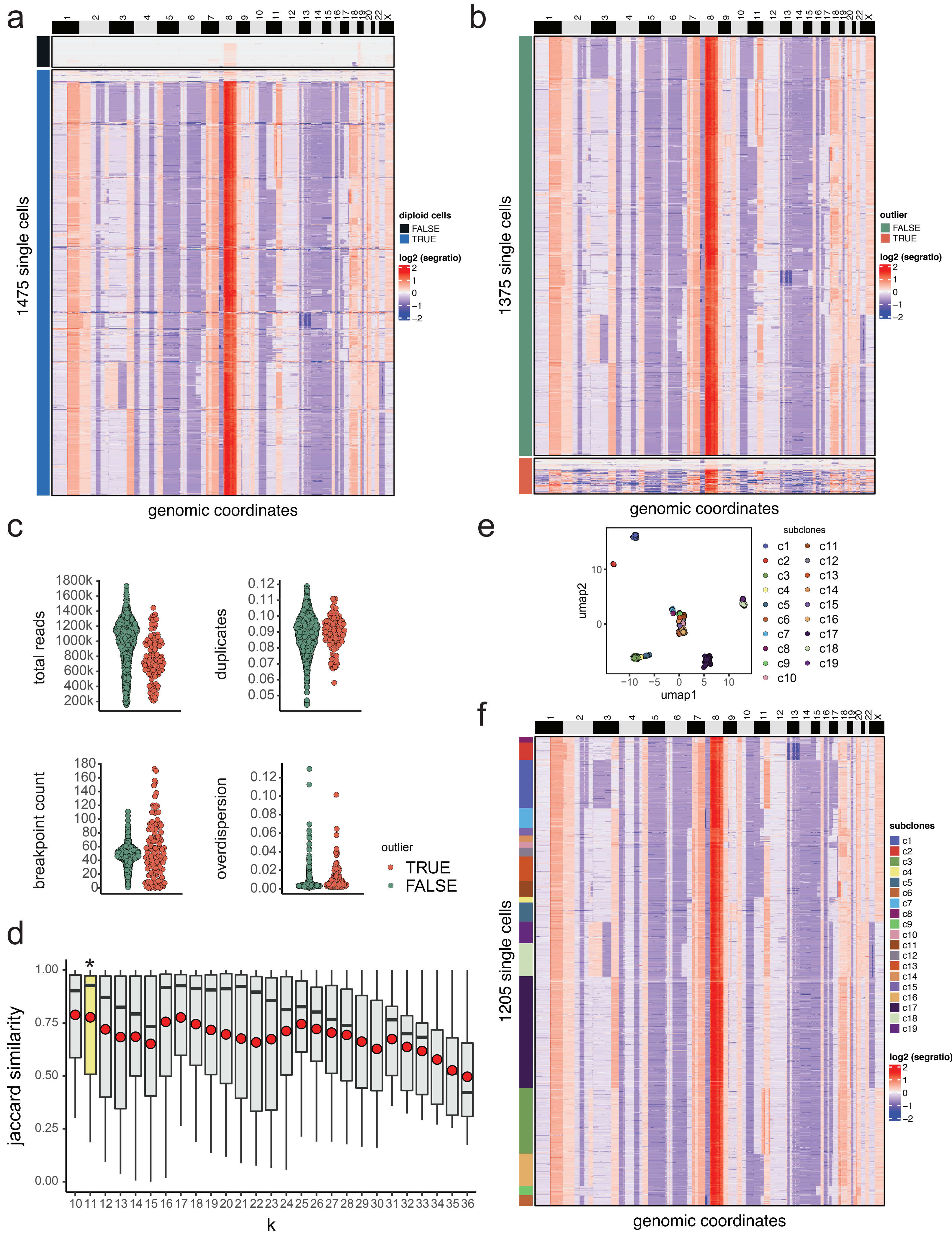
a
b
c
d
e
f
1475 single cells
1375 single cells
1205 single cells
genomic coordinates
diploid cells
FALSE
TRUE
outlier
FALSE
TRUE
log2 (segratio)
total reads
duplicates
breakpoint count
overdispersion
jaccard similarity
k
umap1
umap2
subclones
c1 c2 c3 c4 c5 c6 c7 c8 c9 c10 c11 c12 c13 c14 c15 c16 c17 c18 c19

**Extended Data Fig. 3: CopyKit workflow and clonal substructure of a liver metastasis from a primary breast cancer.**

**a, b,** Clustered heatmap of single cell copy number profiles from the BL2 metastatic liver tumor from a breast cancer patient, in which annotation bar represents classification of cells into diploid or aneuploid (a) or classification into low-quality/high-quality cells (b). **c,** Swarm plots of QC metrics, including the total number of reads and read duplicates (top panels) and overdispersion and counts of breakpoints (bottom panels). **d,** Boxplot distribution of Jaccard Similarity from the clustering parameters grid-search for sample BL2 in which red dots represent the mean Jaccard Similarity and the star indicates the optimal K selected. **e,** reduced dimension embedding UMAP of single cell copy number data, in which colored points represent subclones. **f,** Clustered heatmap of single cell copy number profiles for BL2, in which the annotation bar shows groups of subclone clusters.

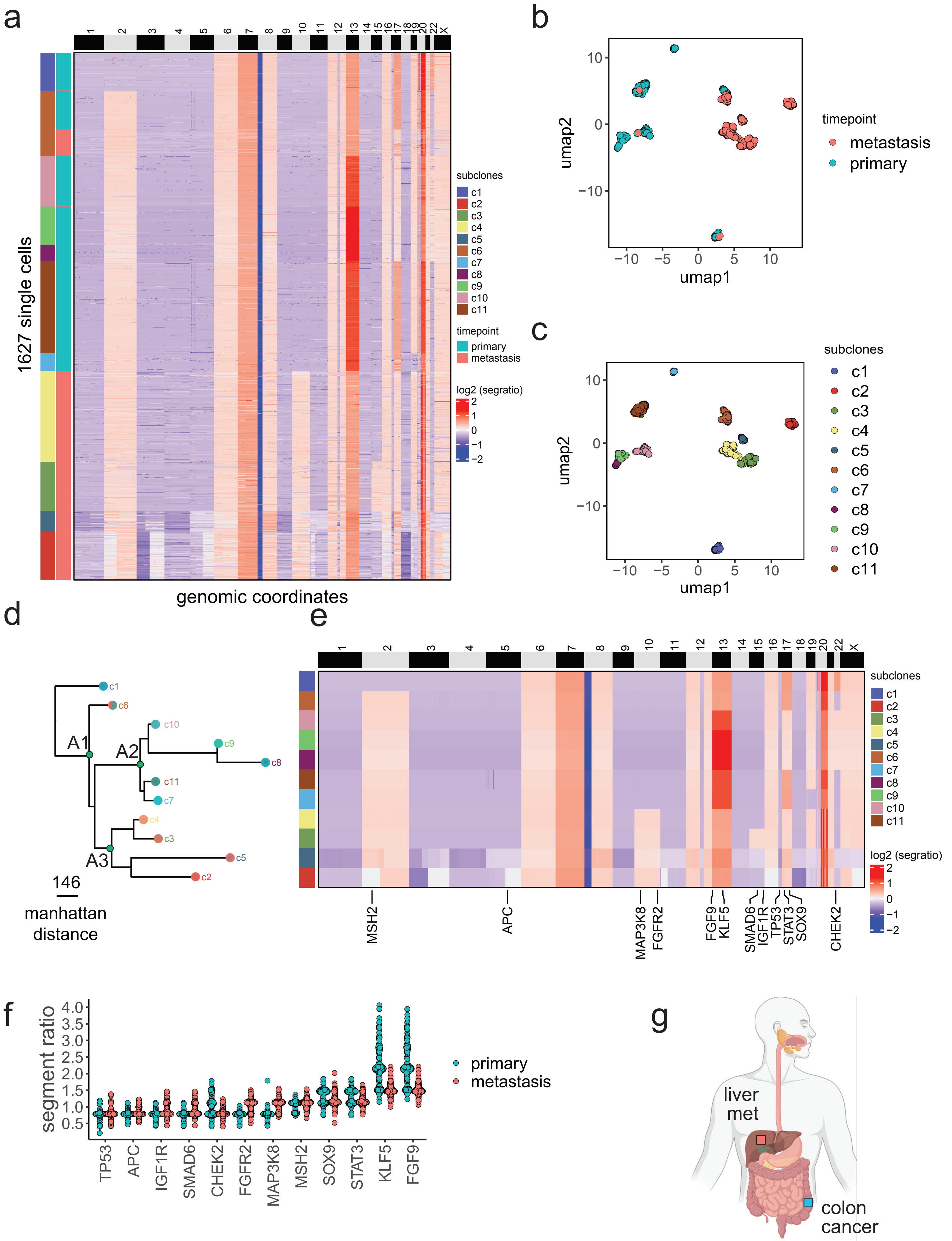

a
b
c
d
e
f
g
1627 single cells
genomic coordinates
1 2 3 4 5 6 7 8 9 10 11 12 13 14 15 16 17 18 19 20 22 X
subclones
c1
c2
c3
c4
c5
c6
c7
c8
c9
c10
c11
timepoint
primary
metastasis
log2 (segratio)
2
1
0
−1
−2
umap1
umap2
10
0
−10
−5
5
timepoint
metastasis
primary
subclones
A1
A2
A3
146
manhattan distance
MSH2
APC
MAP3K8
FGFR2
FGF9
KLF5
SMAD6
IGF1R
TP53
STAT3
SOX9
CHEK2
segment ratio
4.0
3.5
3.0
2.5
2.0
1.5
1.0
0.5
TP53
APC
IGF1R
SMAD6
CHEK2
FGFR2
MAP3K8
MSH2
SOX9
STAT3
KLF5
FGF9
primary
metastasis
liver met
colon cancer

**Extended Data Fig.4: Dissemination from a second patient with a primary colorectal tumor and matched liver metastasis.**

**a,** Clustered heatmap of single cell copy number profiles from a primary colorectal tumor and matched metastatic liver tumor, in which the vertical annotation bar represents tumor site on the right side, and left side annotation bar represents subclones. **b, c,** Reduced dimension embedding UMAP of single cell copy number data, in which colored points represents tumor organ sites (a) and subclones (b). **d,** Minimum evolution tree of the consensus profiles of subclones, in which colored tip labels represent tumor organ sites. **e,** Consensus heatmap of subclones annotated with cancer genes. **f,** Swarm plot showing the distribution of inferred copy number events in which colored points represent the different tumor organ sites. **g,** Anatomical schematic showing the organ sites from where the primary and metastatic tumors tissues were collected from the colon cancer patient.

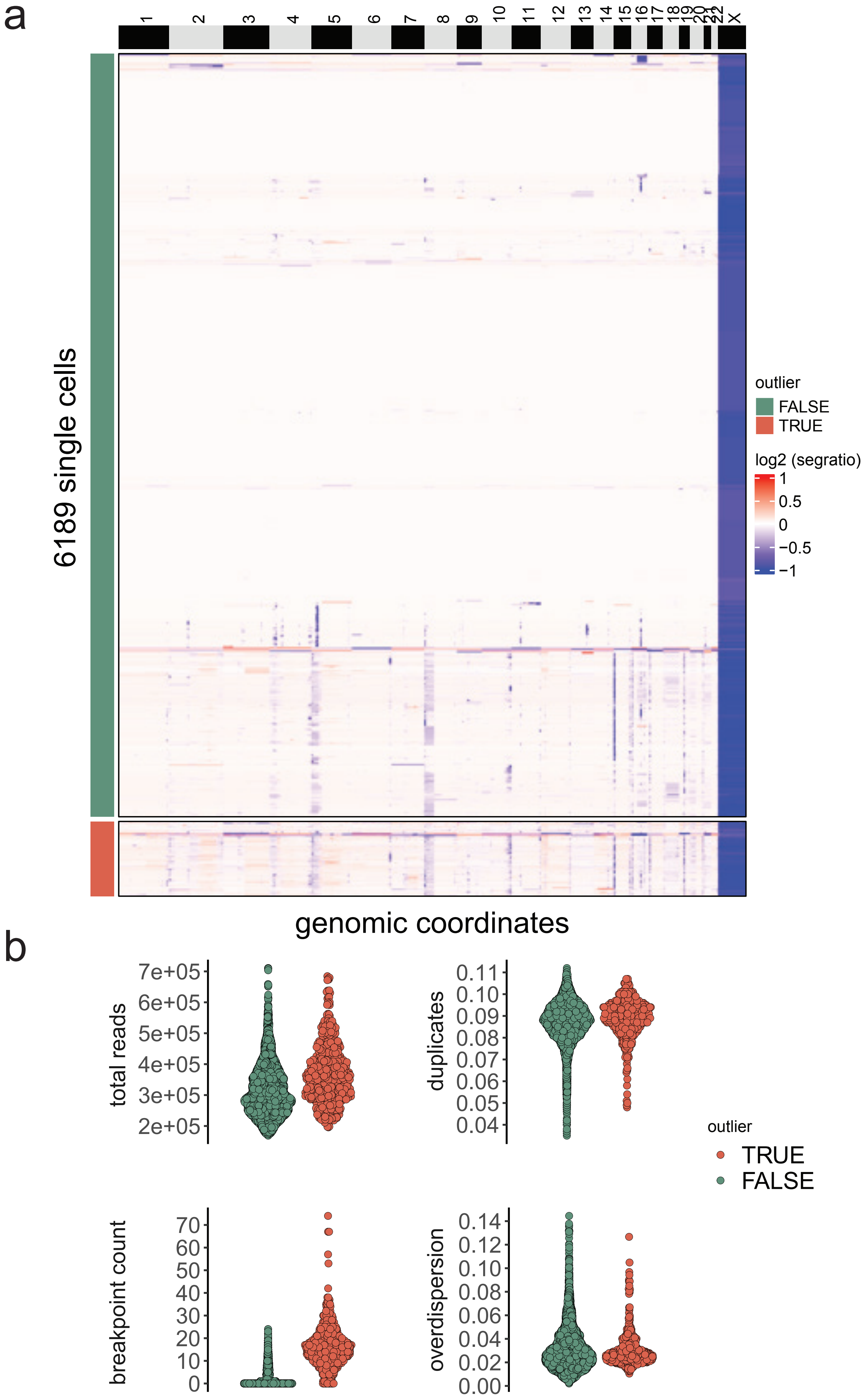
a
X
6189 single cells
outlier
FALSE
TRUE
log2 (segratio)
1
0.5
0
−0.5
−1
genomic coordinates
b
total reads
7e+05
6e+05
5e+05
4e+05
3e+05
2e+05
duplicates
0.11
0.10
0.09
0.08
0.07
0.06
0.05
0.04
outlier
TRUE
FALSE
breakpoint count
70
60
50
40
30
20
10
0
overdispersion
0.14
0.12
0.10
0.08
0.06
0.04
0.02
0.00

**Extended Data Fig. 5 CopyKit workflow applied to a Direct Library Preparation + dataset.** **a,** Heatmap of single cell copy number profiles from the A90533C lymphoblast DLP+ dataset, in which annotation bar represents classification of cells into low-quality (n = 552) or high-quality (n = 5640). **b,** Swarm plots of QC metrics, including total number of reads and read duplicates (top panels) and overdispersion and counts of breakpoints (bottom panels) for the A90533C DLP+ dataset.

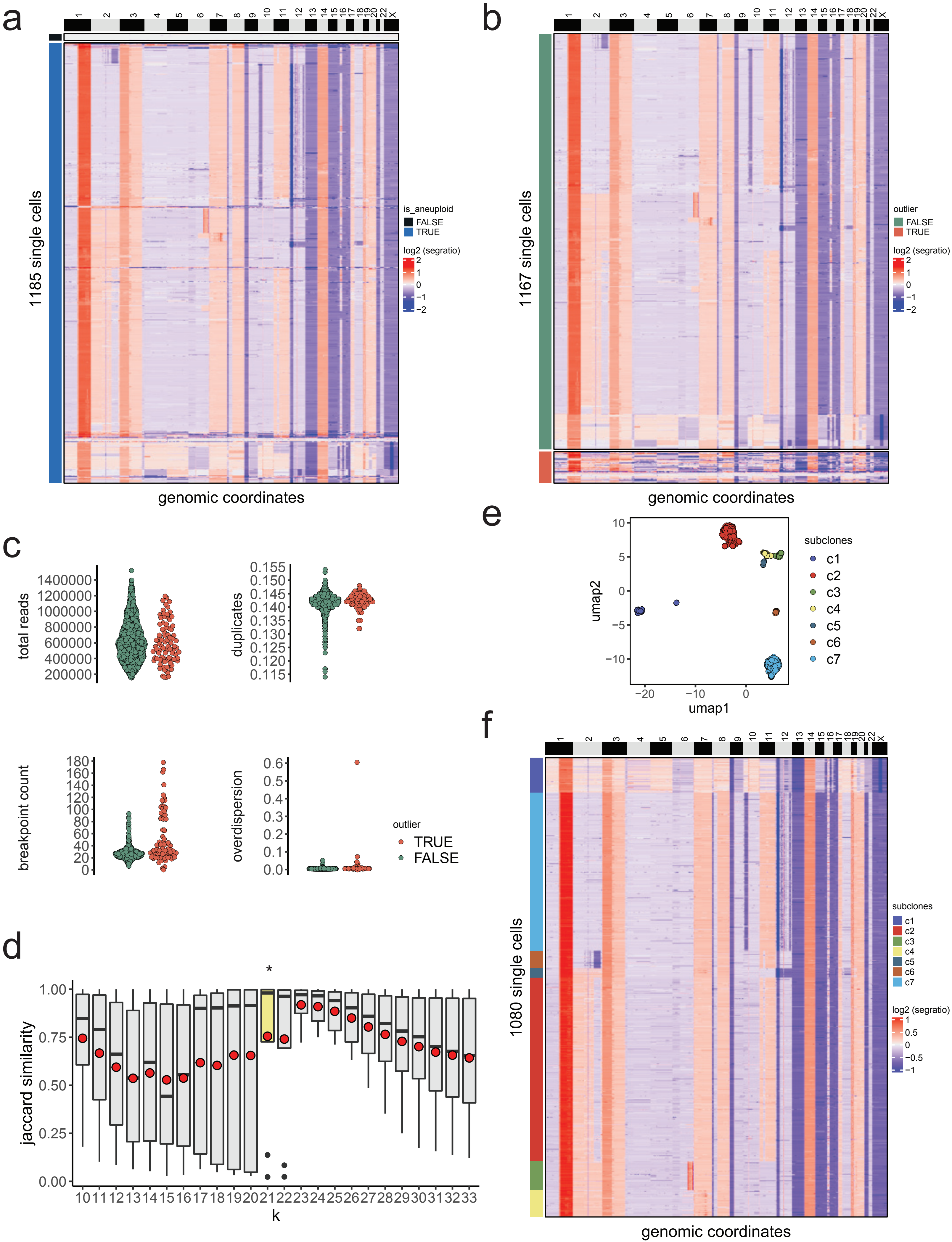
a
b
c
d
e
f
1185 single cells
1167 single cells
1080 single cells
genomic coordinates
is_aneuploid
FALSE
TRUE
log2 (segratio)
outlier
FALSE
TRUE
total reads
duplicates
breakpoint count
overdispersion
TRUE
FALSE
jaccard similarity
k
*
umap1
umap2
subclones
c1
c2
c3
c4
c5
c6
c7

**Extended Data Fig. 6: Application of the CopyKit workflow on a 10X CNV dataset.**
**a,** Heatmap of single cell copy number profiles for the triple-negative breast cancer sample (TN3) in which diploid cells were identified and distinguished from the aneuploid cancer cells. **b,** Heatmap of single cell copy number profiles showing the identification and filtering of low QC cells from the dataset. **c,** Swarm plot of total number of reads and reads duplicates (top) and overdispersion and counts of chromosome breakpoints (bottom), with colors representing the classification of cells by their data quality. **d,** Boxplot distribution of Jaccard Similarity from the clustering parameters grid-search for sample TN3 in which red dots represent the mean Jaccard Similarity and the star indicates the optimal K value selected for clustering. **e,** reduced dimension embedding UMAP and clustering of single cell copy number data in which colored points represents different clusters of subclones. **f,** Clustered heatmap of single cell copy number profiles for TN3, in which annotation bar represents classifications of single cells into subclones.

| ID | age | sex | tissue | ER-primary | PR-primary | HER2-primary | grade | pathology | treatment | DNA ploidy | total cells | detected diploid cells | n cells after filtering | mean reads/cell after filtering | %dup after filtering |
|---|---|---|---|---|---|---|---|---|---|---|---|---|---|---|---|
| **BL1** | 57 | F | liver | pos | neg | neg | 3 | invasive ductal carcinoma | TPH, Denosumab, Herceptin, Arimidex, Kadcyla, Lapatinib | 4.3 | 1502 | 607 | 793 | 1060689 | 0.08 |
| **BL2** | 49 | F | liver | neg | neg | neg | 3 | invasive ductal carcinoma | Chemotherapy NAC, Cediranib, Olaparib, Radiation, Eribulin, Selenexor | 3.4 | 1475 | 100 | 1205 | 992833.3 | 0.09 |
| **BM1** | 44 | F | breast | pos | pos | pos | 3 | invasive ductal carcinoma | AC, paclitaxel, radiation, tamoxifen,arimidex, trastuzumab, fulvestrant, TDM-1 | 3.8 | 442 | 83 | 125 | 1568292 | 0.44 |
| **BM1** | 44 | F | liver | NA | NA | NA | 3 | NA | | 3.8 | 728 | 239 | 238 | 1050535 | 0.07 |
| **BM1** | 44 | F | pleural effusion | NA | NA | NA | 3 | NA | | 3.9 | 351 | 228 | 45 | 1161715 | 0.08 |
| **CM1** | 66 | M | colon | NA | NA | NA | 3 | adenocarcinoma | chemotherapy, AEE788, CPT-11, erbitux, radiation | 3.5 | 2119 | 35 | 1722 | 1120304 | 0.09 |
| **CM1** | 66 | M | liver | NA | NA | NA | 3 | NA | | 3.0/3.76 | 1659 | 13 | 1351 | 969650 | 0.09 |
| **CM2** | 80 | M | colon | NA | NA | NA | 3 | adenocarcinoma | FOLFOX, Avastin, chemotherapy, oxaliplatin | 2.7 | 1063 | 8 | 897 | 1011992 | 0.09 |
| **CM2** | 80 | M | liver | NA | NA | NA | 3 | NA | | 2.7 | 1003 | 6 | 730 | 1098369 | 0.08 |
